# Cajal-Retzius fate specification is disrupted by constitutive activation of β-Catenin in hem progenitors

**DOI:** 10.64898/2026.02.09.704731

**Authors:** Amrita Singh, Arpan Parichha, Debarpita Datta, Mallika Chatterjee, Shubha Tole

## Abstract

Cajal-Retzius cells (CR cells) are the earliest born neurons in the cerebral cortex, and have been implicated in regulating neuronal migration and development of circuitry. A major source of CR cells is the cortical hem, a signaling center at the dorsal telencephalic midline. The hem functions as the hippocampal organizer via canonical WNT signaling and hem progenitors are therefore exposed to high levels of WNT ligands. We tested whether constitutive stabilization of *β-Catenin* (gain of function, GOF) in the mouse cortical hem progenitors supports CR cell production. We find that although neurons are produced from the hem, they do not acquire molecular features of CR cell identity. The trajectory of differentiation examined using single-cell transcriptomics reveals that immature CR cells normally display a *Tbr2+* stage, which is absent upon *β-Catenin* GOF. These data indicate that CR progenitors in the hem are sensitive to levels of stabilized β-Catenin and that a *Tbr2+* stage may be important for the acquisition of CR cell identity.

## Introduction

Cajal-Retzius cells (CR cells), the earliest-born neurons of the mammalian brain, are a transient population that are considered to be essential cellular players in multiple aspects of neocortical development (Bielle et al., 2005; De Frutos et al., 2016; Goffinet et al., 2017; Causeret et al., 2021; Riva et al., 2019; 2023; Genescu et al., 2022; Damilou et al., 2024). Although CR cells were first identified and described more than a century ago (Retzius, 1893; Ramon y Cajal, 1909), the molecular basis of their fate specification is still not well understood.

CR cells that overlie the neocortex and the hippocampus are primarily derived from the cortical hem (Yoshida et al, 2006), a neuroepithelial domain at the medial boundary of the dorsal telencephalon. The hem consists of dividing progenitors that secrete WNT morphogens which induce the hippocampus in adjacent cortical neuroepithelium (Lee et al., 2000; Mangale et al., 2008).

Hem progenitors are themselves exposed to autocrine WNT ligands, yet how canonical WNT signaling operates in hem-derived lineages is an open question. In a previous study, we reported that loss of function of *β-Catenin* in the hem did not affect the specification of CR cells, although it resulted in their mis-migration (Parichha et al., 2022b). Here, we examine the developmental trajectory of CR cells using scRNAseq when canonical WNT signaling is constitutively activated in their progenitors.

Canonical WNT signaling stabilizes cytoplasmic β-Catenin by preventing it from being tagged for degradation. β-Catenin translocates to the nucleus and participates in activation of downstream targets (Hayat et al., 2022). We used a conditional *β-Catenin* gain-of-function (GOF) line (Harada et al., 1999, Parichha et al., 2022a) together with *Lmx1aCre* (Chizhikov et al., 2010) to drive constitutively active β-Catenin in hem progenitors. We report that while this perturbation permits neuronal differentiation, CR cell identity is not acquired in the resulting hem-derived neurons. However, when *β-Catenin* GOF is effected only in postmitotic CR cells, CR cell differentiation proceeds normally. Using scRNAseq, we identify that the *Eomes/Tbr2*+ stage, seen in immature CR cells, is not seen upon *β-Catenin* GOF in hem progenitors. Our results indicate that unrestrained canonical WNT signaling in hem progenitors prevents a key transition through a *Tbr2*+ stage, therefore, progenitors must limit their response to canonical WNT signaling in order to produce bonafide CR cells.

## Results

### Cortical hem progenitors giving rise to Cajal Retzius cells downregulate canonical WNT signaling during the course of differentiation

*Lmx1a* expression is specific to the hem and its derivatives within the telencephalic midline. We used an *Lmx1aCre* mouse line (Chizhikov et al., 2010) which faithfully marks lineages arising from the hem when crossed with an Ai9 reporter (Fig. 1A, Chizhikov et al., 2010; Parichha et al., 2022a).

**Figure 1:**
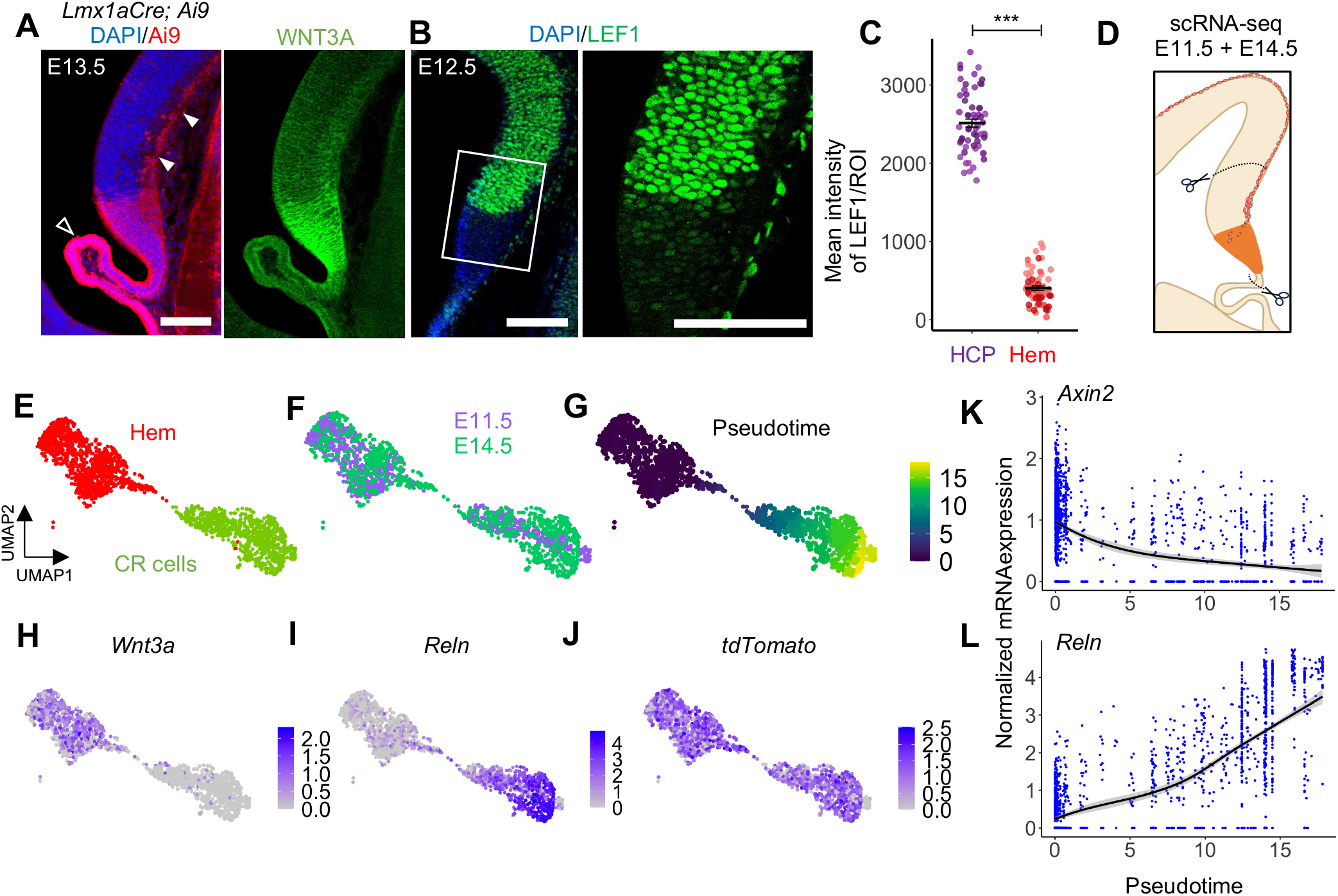
Canonical WNT signaling in the cortical hem and the CR cell lineage. (A) The Ai9 reporter driven by *Lmx1aCre* is detected the cortical hem and its derivatives (arrowheads, CR cells; open arrowhead, choroid plexus epithelium) in an E13.5 *Lmx1aCre*; *Ai9* brain, and WNT3A protein is detected in the hem. (B, C) Nuclear LEF1 staining (B) is low in the hem compared to adjacent hippocampal tissue, quantified in progenitor cells near the VZ (C). (D) Schematic showing tissue isolated for scRNA-seq. (E-J) UMAPs representing tdTomato+ cells from E11.5 and E14.5, color-coded by cell type (E); by age (F); by pseudotime (G); showing the expression of *Wnt3a, Reln* and *tdTomato* (H-J). (K, L) Pseudotime plots showing the levels of *Axin2* and *Reln* across the CR cell differentiation axis. Scatterplot in (C) displays Mean ± SEM. Statistical test (C): Shapiro-Wilk normality test, followed by Welch’s two-sample t-test; p < 0.0001; *p < 0.05; **p < 0.01; ***p < 0.001; ns if p value > 0.05. Statistical analysis performed on replicate-level means. For (A-C), N ≥ 3 brains (biologically independent replicates); (C), N=3 (biologically independent replicates), n = 20 cells each per location, per biological replicate. Scale bars: 100 μm (all images in A and B).

Even though the hem expresses WNT ligands, LEF1, a downstream effector and target of canonical WNT signaling, is detected at significantly lower levels in hem progenitors as compared to the adjacent hippocampus, suggesting that they may be sensitive to the level of canonical WNT activation they experience. (Fig. 1B, C)

We profiled the developing telencephalic midline at ages E11.5 and E14.5 when newborn hem-derived CR cells populate the dorsal telencephalon (Hevner et al, 2003). The hem and a portion of the adjacent hippocampal primordium was isolated from wild-type *Lmx1aCre*; *Ai9* brains and scRNA-seq libraries were prepared. To identify cells arising from the hem, a region corresponding to the tdTomato fluorescent reporter protein was added to the mm10 genome assembly, which was used as a reference for alignment. After quality control, filtering and integration, a total of 5758 cells from E11.5 and 6252 cells from E14.5 were retained, which were projected on low-dimensional UMAP spaces for visualization purposes. (Fig. S1A-B). Louvain clustering and annotation based on known marker genes (Fig. S1C-D) identified clusters corresponding to the hem and its derivatives, the CR cells and the choroid plexus epithelium, which were validated by tdTomato expression, subsetted and projected on a new landscape. *Ttr* and *Htr2c* expression identified choroid plexus cells which were removed. The remaining 1881 (hem+CR) cells (Fig. 1E, F) were used to derive a pseudotime trajectory of differentiation from hem cells to CR cells (Fig. G), using the Monocle3 package (Trapnell et al, 2014). As expected, the expression of *Reln* rose along the pseudotime axis (Fig. 1L). However, the expression of *Axin2*, a well-known canonical WNT signaling target gene, decreased (Fig. 1K), suggesting that canonical WNT signaling is downregulated in the course of CR cell differentiation. This raised the question of whether this downregulation is necessary for CR cell maturation.

### β-Catenin stabilization in the cortical hem compromises the Cajal-Retzius cell fate

To test the effect of constitutive activation of canonical WNT signaling on CR development, we used an *Lmx1aCre* driver together with a well-described *β-Catenin* GOF mouse line (Harada et al., 1999, Parichha et al., 2022a). In this line, Cre-mediated recombination leads to the deletion of exon 3 of the *β-Catenin (Ctnnb1)* gene. This region contains sites at which β-Catenin is phosphorylated by GSK3β, typically leading to its degradation. The resultant recombined *β-Catenin* allele produces constitutively active and functional β-Catenin (Harada et al., 1999). In control brains, immunohistochemistry for Reelin, TRP73 labeled CR cells that also displayed the Ai9 reporter. In contrast, *Lmx1aCre; β-Catenin* GOF brains displayed minimal staining for any of these CR cell markers, or GRM2 (Causeret et al., 2021), a marker of differentiated CR cells, even though Ai9+ cells accumulated in the marginal zone similar to controls (Fig. 2A-C, Fig. S2A, D). Consistent with the immunohistochemistry, qPCR for several CR cell markers (*Reln, Trp73, Lhx5*, and *Ppp2r2c*) from microdissected control and *Lmx1aCre; β-Catenin* GOF medial tissue revealed a significant downregulation upon hem-specific *β-Catenin* GOF (Fig. 2D).

**Figure 2:**
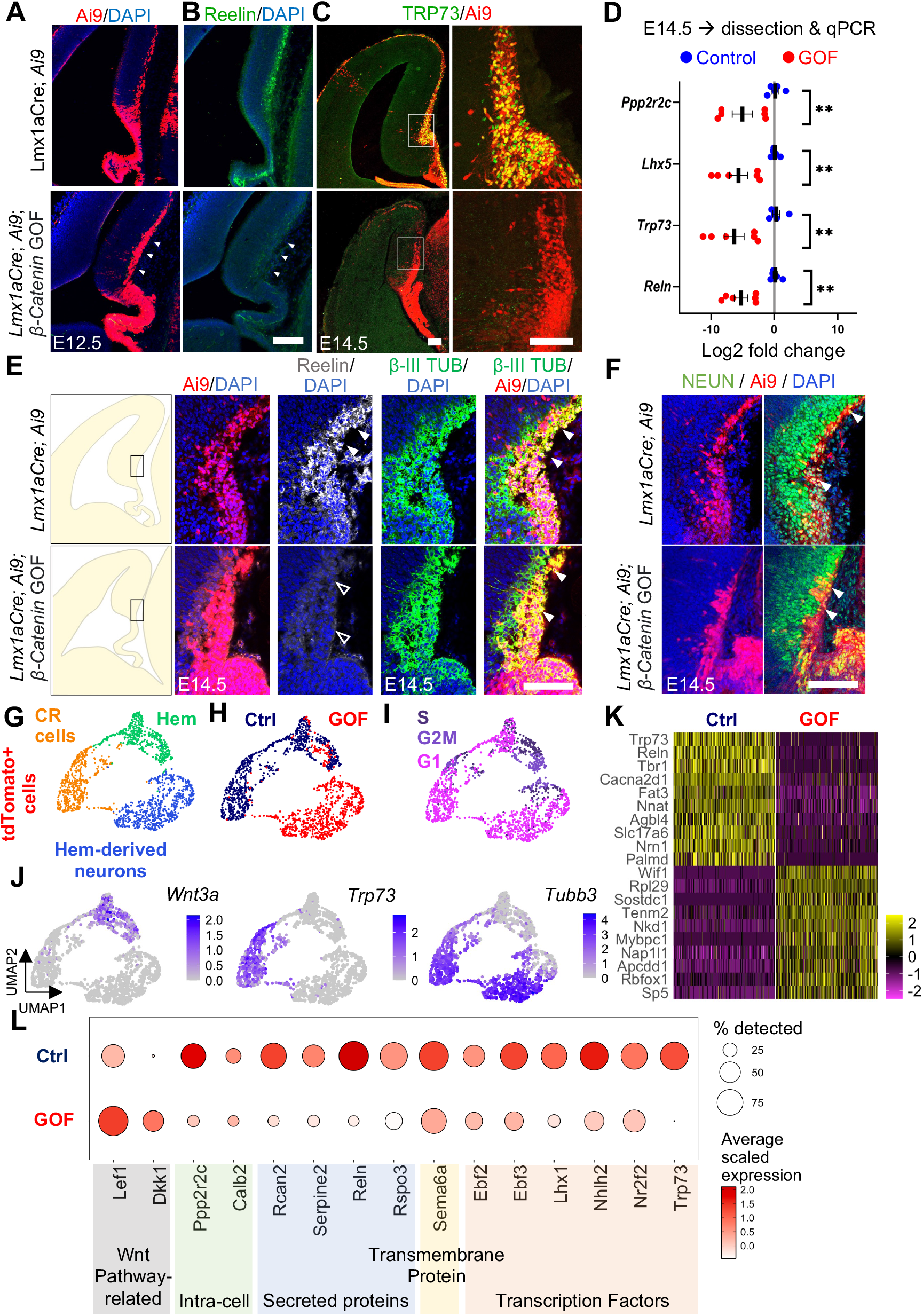
CR markers are undetectable in hem-derived neurons in β-CATENIN GOF brains. (A) Ai9 is seen in the cortical hem and its derivatives in an E12.5 control and *Lmx1aCre; β-Catenin* GOF brain. (B) Reelin staining in the same section as (A). (C) TRP73 staining at E14.5 co-localizes with Ai9 in the control but is undetectable in the *β-Catenin* GOF brain. (D) Genes enriched in CR cells are downregulated in midline tissue *β-Catenin* GOF brains at E14.5. (E, F) Reelin, βIII-Tubulin and NEUN staining is seen in Ai9+ cells in control brains (arrowheads). In *β-catenin* GOF brains there is no detectable Reelin (open arrowheads) (E) but βIII-TUBULIN and NEUN staining is seen in Ai9+ cells. (G, J) UMAPs representing tdTomato+ cells from E14.5 control and *β-catenin* GOF midline, color-coded by cell type (G); by age (H); by genotype (I); showing the expression of *Wnt3a, Reln* and *tdTomato* (J). (H) Heatmap of scaled expression of top 10 differentially expressed genes in control and *β-Catenin* GOF neurons. (I) Dot plots showing scaled expression levels of CR cell enriched genes in control and *β-Catenin* GOF neurons. Scatterplot in (D) displays Mean ± SEM. Statistical test(D): Multiple Mann-Whitney Tests; p < 0.0001; *p < 0.05; **p < 0.01; ***p < 0.001; ns if p value > 0.05. For (A-C, E-F), N ≥ 3 brains (biologically independent replicates); for (D) N=6 (control), 7 (GOF) biologically independent replicates. Scale bars: 100 μm (all images in A, B, C, E and F).

Surprisingly, the hem-derived Ai9+ cells were positive for β-III-Tubulin and NEUN (RBFOX3), which confirmed their neuronal identity (Fig. 2E-F). We tested whether there was a delay in CR maturation by examining *Reelin* expression using *in situ* hybridization at a range of time points (E12.5–E18.5) when CR cells are seen covering the dorsal telencephalon. In *β-Catenin* GOF brains, *Reelin* expressing cells were almost undetectable in the medial telencephalon (black arrowheads, Fig. S2B), although CR cells generated at the pallial-subpallial boundary (PSB), did express *Reelin* (red arrowheads, Fig S2B). This provided a useful internal control since the PSB does not express *Lmx1a* and therefore was unaffected by the *β-Catenin* GOF mutation.

An alternate possibility that may explain the lack of CR cells in the *β-Catenin* GOF brains is cell death. Ai9+ cells in the marginal zone of the *β-Catenin* GOF brains showed no increase in TUNEL+ cells at E12.5 (Fig. S2C), however. CR cells are the only known glutamatergic cortical neuron type that do not express FOXG1, which is considered to be a repressor of CR fate (Hanashima et al., 2004; Elorriaga et al., 2023). However, no upregulation of FOXG1 was seen in the hem-derived Ai9+ cells in the marginal zone of the *β-Catenin* GOF brains (Fig. S2E).

In summary, these data indicate that when β-CATENIN is stabilized in the cortical hem progenitors, CR cell differentiation is compromised, although aspects of neuronal differentiation are preserved.

To examine how CR cell differentiation is compromised upon *β-Catenin* GOF, we performed scRNA-seq on E14.5 midline tissue of the *Lmx1aCre β-Catenin* GOF brains and compared it with E14.5 control data (Fig. 1). After quality control and filtering, a total of 11216 cells from control and GOF samples were merged and projected on the same landscape for visualization purposes (Fig. S3A-C). As before, Ai9+ clusters were subsetted, choroid plexus epithelium cells removed, and the remaining Ai9+ cells were subjected to further analysis.

Three broad clusters were identified: hem progenitors, identified by *Wnt3a*, CR neurons, identified by *Trp73*, and a cluster that displayed neuronal marker *Tubb3* (*β-III tubulin*) but neither *Wnt3a* nor *Trp73*. We refer to this cluster as “hem-derived neurons”. Both control and *β-Catenin* GOF cells were found in the hem cluster. The CR cluster contained largely control cells and only a few *β-Catenin GOF* cells, whereas the third cluster consisted exclusively of *β-Catenin* GOF cells. This population of neurons in the mutant, that failed to achieve bonafide CR identity, was analyzed further.

Differential gene expression analysis between the CR cells cluster and the hem-derived neurons cluster revealed a total of 2053 genes upregulated in the CR cells and 1872 genes upregulated in the hem-derived neurons (Fig. S3D) of which the top 15 in each condition are shown in a heatmap (Fig. 2K). As expected, targets of the canonical WNT signaling pathway, *Lef1* and *Dkk1* were upregulated in the GOF cells compared to the controls, thus indicating that this pathway was indeed transcriptionally active at a higher level in the GOF cells (Fig. 2L). Consistent with the qPCR and immunohistochemistry results, well-established molecular markers of CR cells (Causeret et al., 2021), were downregulated in the GOF hem-derived neurons compared with controls (Fig. 2L). Multiple sources of CR cells produce molecularly distinct subtypes of this population. Therefore, we used markers comprehensively characterised in Moreau et al., 2021 to examine whether the GOF cells had switched identity to a different CR subtype, including midline-originating CR populations from the septum or thalamic eminence (Fig. S3E). However, the GOF hem-derived neurons did not display markers consistent with a switch to a different CR subtype.

### Investigating the identity of the transformed neurons in the β-Catenin GOF

We further investigated whether the *β-Catenin* GOF hem-derived neurons may have differentiated to a completely different neuronal subtype in the brain. We curated a list of well-established marker genes known to be expressed in the neocortex, the hippocampus, and in cortical interneurons. (Tole et al., 1997; 2001; Muralidharan et al., 2017; Molyneaux et al., 2007, Ba et al., 2023). A comparison of the scaled expression of these genes between the control and GOF neurons reveals a reduction in layer 1 markers *Lhx5* and *Reln*, which is consistent with a loss of CR identity. However, none of the other neuronal subtype markers appear to change (Fig. 3A).

**Figure 3:**
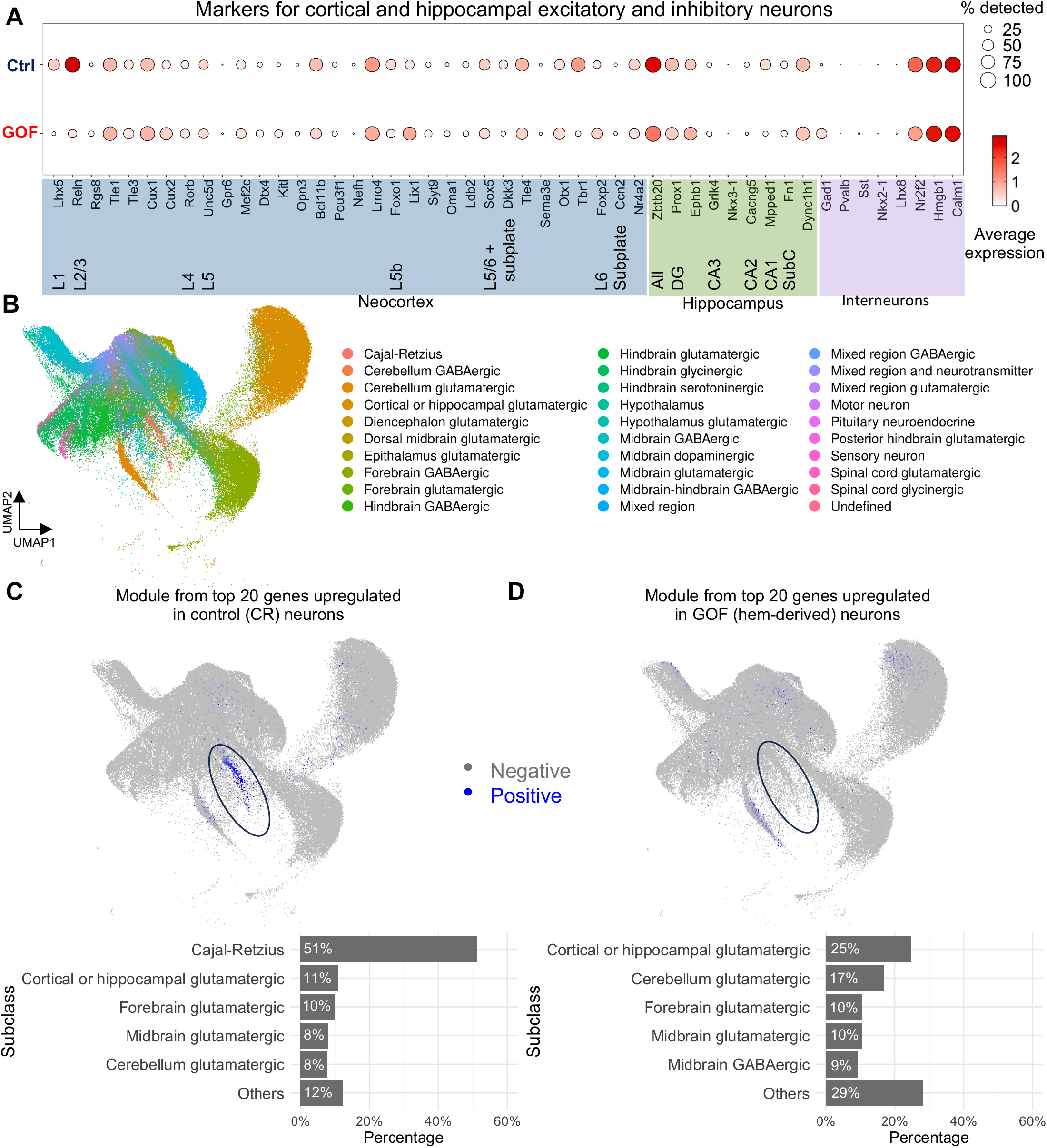
Hem-derived neurons from *β-Catenin* GOF brains do not map to any known category of neurons in the brain. (A) Dot plots showing scaled expression levels of neocortical and hippocampal excitatory and inhibitory markers in control and *β-Catenin* GOF neurons. (B) UMAP showing the “neuron” class subsetted from the La Manno 2021 dataset, colour-coded by subclass (C-D) UMAP from (B), in which cells above the 99th percentile of module scores for genes upregulated in control (oval, C) and *β-Catenin* GOF (oval, D) neurons are represented in blue. Bar graphs show the top 5 subclasses of cells above 99th percentile of the module scores.

To investigate their identity in an unbiased manner, we used a publicly available scRNA-seq dataset from the developing mouse brain (La Manno et al., 2021). We hypothesized that if the hem-derived *β-Catenin* GOF neurons are immature, a developing mouse brain dataset would be better suited for analysis. We subsetted the La Manno dataset to cells annotated as neurons, which retained a total of 111063 cells divided into 30 subclasses (Fig. 3B). We used the AddModuleScore() function in the Seurat package to create a module “signature” (Supplementary Table 1) for control CR cells and hem-derived *β-Catenin* GOF neurons to enable mapping to this curated dataset of 30 neuronal subclasses. Fig. 3C displays neurons in the La Manno dataset that matched to a module score above the 99th percentile for the control CR module, a large fraction of which (51.25%), as expected, are in the cluster annotated as CR cells in the original dataset. Fig. 3D displays neurons in the La Manno dataset that matched to a module score above the 99th percentile for the hem-derived *β-Catenin* GOF neurons. As evident from the UMAP, these cells do not map to any annotated cluster preferentially, but instead are distributed as “cortical or hippocampal glutamatergic” or other glutamatergic neurons, with a small fraction mapping to “midbrain GABAergic neurons” (Fig. 3D). Together, these results suggest a mixed neuronal identity arising from *β-Catenin* GOF in the hem.

### Gene expression dynamics are disrupted in hem-derived *β-Catenin* GOF neurons

To explore the gene expression dynamics in hem-derived *β-Catenin* GOF neurons, Monocle3 was used to learn trajectories separately in the control and GOF cells. The “hem” cluster was assigned as the root cells in each case (Fig. 4A). We examined a set of genes known to be sequentially expressed during cortical neurogenesis along a pseudo-differentiation axis, including *Pax6, Eomes* and *Tbr1* (Englund et al., 2005). This revealed a profoundly disrupted developmental trajectory in the GOF neurons (Fig. 4B). Only *Pax6* displayed a somewhat comparable trajectory in GOF neurons and controls. *Neurog2, NeuroD2*, and *Tbr1* dynamics indicated the GOF neurons eventually appeared to achieve comparable expression to controls, but displayed disrupted trajectories. Unexpectedly, the GOF neurons did not appear to express *Eomes/Tbr2* at all, in contrast to control neurons that displayed a peak at an early stage in the pseudo-differentiation timeline (Fig. 4B). We examined whether this apparent loss of an *Eomes/Tbr2*+ stage was also evident using Immunohistochemistry. At E12.5, the number of Ai9+ TBR2+ cells was greatly reduced in *Lmx1aCre; β-Catenin* GOF brains compared to controls (arrowheads, Fig. 4C).

**Figure 4:**
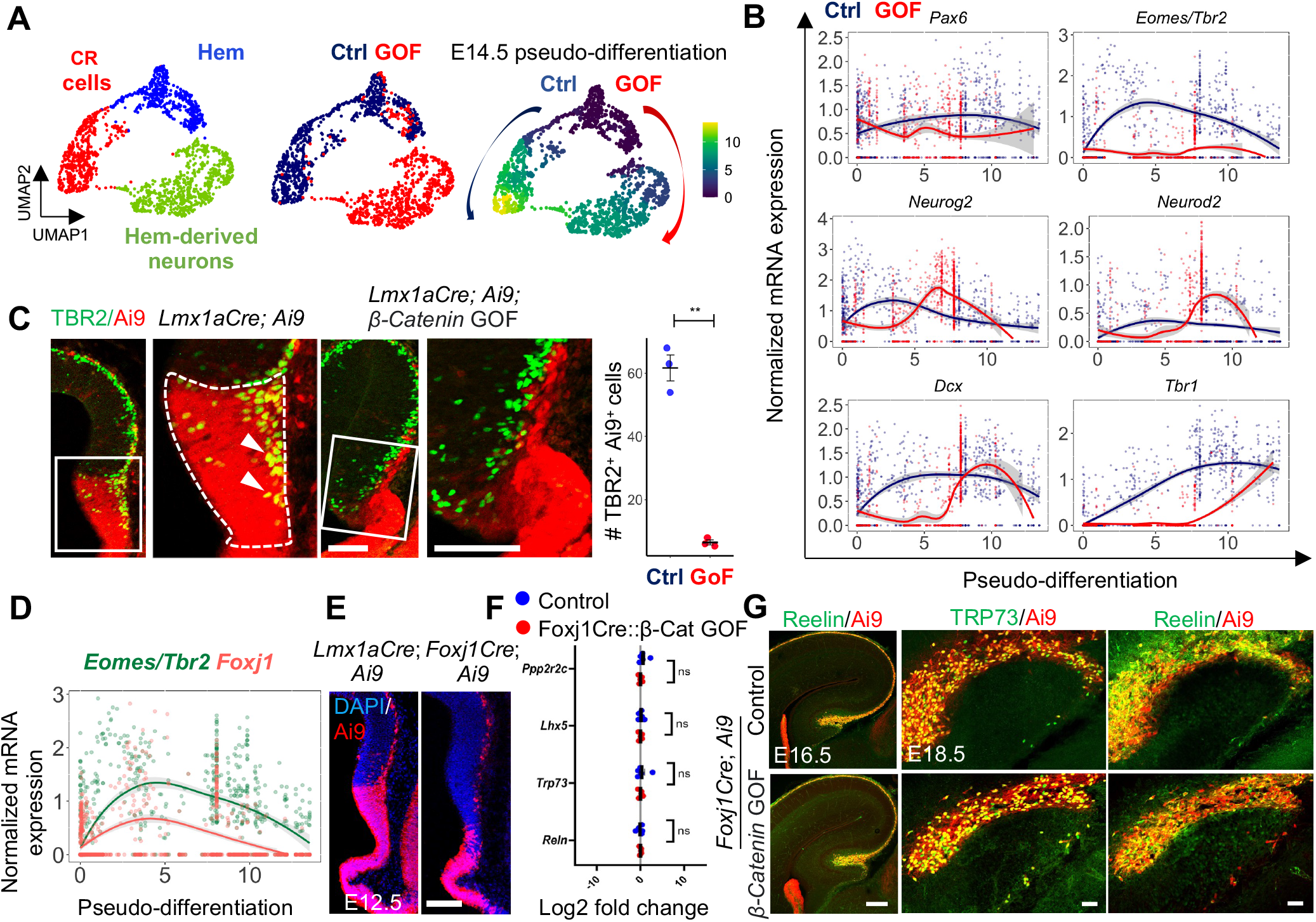
Disrupted developmental trajectory of hem-derived neurons from β-Catenin GOF brains. (A) UMAPs from Figure 2 representing tdTomato+ cells from control and *β-Catenin* GOF, color-coded by cell type; by genotype; and by pseudo-differentiation trajectory derived from Monocle3. (B) Normalized mRNA expression of *Pax6, Eomes/Tbr1, Neurog2, Neurod2, Dcx* and *Tbr1* across the pseudo-differentiation axis for control (blue) and *β-Catenin* GOF (red). Thick lines represent Loess smoothed curves. (C) TBR2 staining is seen in Ai9+ cells in controls (arrowheads) but not in *β-Catenin* GOF brains. Dashed lines mark the ROIs in which TBR2+Ai9+ cells were quantified. (D) The *Eomes* and *Foxj1* expression trajectory along the pseudo-differentiation axis in control neurons (E) Ai9 reporter expression at E12.5 comparing *Lmx1a*Cre and *Foxj1*Cre activity. *Foxj1*Cre is not active in hem progenitors but is seen in CR cells and the choroid plexus epithelium. (F) Genes enriched in CR cells are unchanged in midline tissue of *Foxj1Cre; β-Catenin* GOF brains at E14.5. (G) TRP73 and REELIN staining co-localizes with Ai9+ cells in the hippocampal fissure in both control and *Foxj1Cre*; *β-Catenin* GOF brains at E16.5 and E18.5. Scatterplots in (C) and (F) display Mean ± SEM. Statistical test (C): Shapiro-Wilk normality test, followed by Welch’s two sample t-test, (F) Multiple Mann-Whitney Tests; p < 0.0001; *p < 0.05; **p < 0.01; ***p < 0.001; ns if p value > 0.05. For (C), N=3 (biologically independent replicates), (E), N=5 (biologically independent replicates), (F), N=6 (biologically independent replicates), (G), N=3 (biologically independent replicates). Scale bars: 100 μm (all images in C, E and G).

Interestingly, in Hodge et al., 2013, when *Tbr2* itself was disrupted in CR cells using *NestinCre*, p73 expressing cells were seen in the hem. However, the recombination was performed with *NestinCre* which acts from E11.5, much later than *Lmx1aCre* (Chizhikov et al., 2010).

### Stabilization of β-Catenin at an intermediate stage of CR cell differentiation does not alter their fate

A recent study identified a multi-ciliation gene module to be critical after the apical progenitor stage for CR cell differentiation (Moreau et al., 2023). We examined the gene expression dynamics of *Foxj1*, a well-established marker of multi-ciliated cells, in the control E14.5 CR cell dataset, and found it to parallel *Eomes/Tbr2* trajectory (Fig. 4D). We therefore tested the effect of stabilization of β-Catenin using *Foxj1Cre* (Zhang et al., 2007). First, we examined Ai9+ cells using *Foxj1Cre* and confirmed that both CR cells and choroid plexus epithelium cells were positive as expected but in contrast to *Lmx1aCre*, the hem progenitors were not Ai9+ (Parichha et al., 2022a, Fig. 4E). First, we examined CR cell marker genes *Reelin, Trp73, Lhx5, Chst8*, and *Ppp2r2c* using qPCR of microdissected control and *Foxj1Cre; β-Catenin* GOF medial tissue. In contrast to the findings in *Lmx1aCre; β-Catenin* GOF brains (Fig. 2D), none of these genes displayed any change from controls in *Foxj1Cre; β-Catenin* GOF brains (Fig. 4F). Immunohistochemistry for Reelin and TRP73 revealed CR cells occupying the hippocampal fissure in a manner indistinguishable from controls at E16.5 and E18.5 (Fig. 4G).

Taken together, these data suggest that the CR progenitors in the hem are sensitive to excessive levels of β-Catenin only at an early stage. The *β-Catenin* GOF driven by *Lmx1aCre* in progenitors causes the cells to bypass an *Eomes/Tbr2*+ step and prevents them from acquiring the molecular characteristics of bonafide CR cells. By the time the cells express *Foxj1*, excessive levels of β-Catenin do not disrupt CR differentiation.

In the cortical primordium, increase in β-Catenin levels decreases the number of TBR2+ intermediate progenitors arising from cortical progenitors (Mutch et al., 2009, Mutch et al., 2010). Our study reveals that β-Catenin signaling must be suppressed in hem progenitors in order to ensure CR genesis. Surprisingly, increased β-Catenin in post-mitotic CR cells does not suppress their fate, possibly due to differences in interactions with progenitor-specific transcriptional repressors or altered chromatin accessibility between these two cell types, suggesting an avenue for future studies.

A recent study found that a multiciliation gene module controlled by *Gmnc* is critical for proper CR cell specification without which there is premature cell death (Moreau et al., 2023). Our findings add another significant milestone, that of a *Tbr2+* state, without which neuronal differentiation proceeds, but molecular signatures of CR cells fail to be acquired. Consistent with our finding, when *Tbr2* is disrupted from E12.5 using a *Tbr2CreERT2* driver, there is a loss of Reelin+ cells in the cortex (Mihalas et al., 2016).

In our study, this step seems to be missed, which implies that neurogenesis and fate specification in CR cells may be uncoupled from each other. This may provide a general model for how neurons are produced and acquire their identities.

## Limitations

Two important aspects of CR cell character is their distinctive pattern of tangential migration along the marginal zone of the cortex, as well as their transient nature, since they undergo apoptosis in the second week of postnatal mouse cortex (Ledonne et al, 2016). Examining these angles in the hem-derived neuronal population requires further study.

## Materials and Methods

### Animal strains

The Institutional Animal Ethics Committee of the Tata Institute of Fundamental Research, Mumbai, India has approved all animal protocols used in this manuscript. For the kind gifts of mouse lines used in this study, we thank Kathy Millen (Centre for Integrative Brain Research, Seattle Children’s Research Institute - Lmx1aCre line described in Chizikov and Millen, 2004), Michael J. Holtzman and Yong Zhang (University of Washington, St. Louis - Foxj1Cre line), and Makoto M. Taketo (Kyoto University, Kyoto - *Ctnnb1* exon 3 floxed mouse line). Ai9 reporter mouse line (Stock No. 007909) was obtained from JAX labs. Ambient temperature and humidity conditions were maintained for all mouse lines, with a strict 12hr light-dark cycle. Food and water were available ad libitum. Noon of the day when vaginal plug was observed, was considered embryonic day 0.5 (E0.5). Samples were analyzed from animals of both sexes. Controls are littermates unless stated otherwise. Since germline recombination occurs in the testes of the Lmx1aCre and Foxj1Cre lines, breeding colonies were maintained with the Cre transgene on the female breeders.

Primer sets used in genotyping:

Cre: (Forward) 5’ATTTGCCTGCATTACCGGTC3’, (Reverse) 5’ATCAACGTTTTCTTTTCGG3; 350bp band observed in Cre-positive animals.

β-CATENIN GOF (Exon 3 floxed): (Forward) 5ʹ GCTGCGTGGACAATGGCTAC3ʹ; (Reverse) 5ʹGCTTTTCTGTCCGGCTCCAT3ʹ; 550p band for conditional allele and 350bp band for wild type allele.

### Tissue sections and immunohistochemistry

Mouse embryo dissections were performed in ice-cold PBS. Brains were fixed in 4% PFA overnight at 4°C and equilibrated in 30% sucrose in PBS before sectioning at 25μm thickness. Coronal sections were transferred to a slide mailer (catalogue number EMS 71549-08) and washed with PBS + 0.01% TritonX-100 for 10 minutes, followed by a 2 5-minute washes with PBS + 0.03% TritonX-100. Antigen retrieval was performed by boiling sections in a 10mM sodium citrate buffer (ph = 6) at 90°C for 10 minutes in a water bath. After cooling to room temperature, slides were washed with PBS + 0.01% TritonX-100 for 10 min. Blocking solution of 5% horse serum/Lamb serum in PBS + 0.1% TritonX-100 was added to the slides and they were incubated for 1 hr in a humidified box. This was followed by overnight incubation with primary antibody at 4°C. The next day, they were incubated with secondary antibody at room temperature in a humidified box covered with aluminium foil (to prevent bleaching of the fluorophore). After two washes with PBS and a 10 minute incubation with DAPI, slides were mounted using Fluoroshield mounting medium (Sigma catalogue #F6182). Primary antibodies used: Lef1(rabbit, 1:200, CST catalogue #C12A5), β-CATENIN (Mouse, 1:200, BDbiosciences catalogue #610153), β-CATENIN (Rabbit, 1:50, CST catalogue # 8814), RFP (rabbit, 1:200, Abcam catalogue #ab62341), RFP (Mouse, 1:200, Invitrogen catalogue #MA5-15257), β-III TUBULIN (mouse, 1:100, Promega catalogue #G7128), TRP73 (Rabbit, 1:200, CST catalogue #14620S), REELIN (Mouse, 1:200, Millipore catalogue #MAb5364), NEUN (Rabbit, 1:200, invitrogen catalogue #702022). Secondary antibodies used: Goat Anti Rabbit Alexa fluor 488 (1:200, Invitrogen catalogue #A11034), Goat Anti mouse Alexa fluor 594 (1:200, Invitrogen catalogue #R37121), Goat Anti Rabbit Alexa fluor 568 (1:200, Invitrogen catalogue #A11011), Donkey Anti rabbit Alexa fluor 647 (1:200, Invitrogen catalogue #A31573), Goat Anti Mouse Alexa fluor 647 (1:200, Invitrogen catalogue #A21236), Goat Anti Mouse Alexa fluor 488 (1:200, Invitrogen catalogue #A28175), Goat Anti Rat Alexa fluor 488 (1:200, Invitrogen catalogue #A11006).

### In situ hybridization

A freezing microtome (Leica SM2000R Sliding Microtome) was used to section fixed mouse brains at a thickness of 25μm. Sections were then mounted on superfrost plus slides (Catalogue #EMS 71869-11), post-fixed with 4% (w/v) paraformaldehyde for 15 min, and washed thrice with PBS. The sections were then treated with proteinase K dissolved in Tris-EDTA buffer (1 μg/ml). They were then post-fixed in 4% PFA for 15 min, then washed thrice in PBS. Hybridization was performed for 16 hrs at 70 °C in a buffer with 50%(v/v) formamaide, 2X SSC, and 1%(w/v) SDS; with Digoxigenin (DIG) labelled cRNA probes prepared from the respective plasmids using in vitro transcription. Three stringent washes (45 mins each) in solution X (50% formamide, 2X SSC, and 1% SDS) were followed by a wash with 2XSSC and then 0.2XSSC. Blocking was performed in 10% horse serum in TBST (Tris-buffered saline pH 7.5 with 0.1% Tween-20) for 1 hr. Sections were incubated in alkaline phosphatase-conjugated anti-DIG (Digoxygenin) antibody Fab fragments (1:5000; Roche, catalog #12486523) at 1:5000 in the blocking buffer and incubated at 4 °C overnight. NBT/BCIP substrate (Roche, 4-nitroblue tetrazolium chloride, catalog #70210625; 5-Bromo-4-chloro-3-indolyl phosphate, catalog #70251721) was used for color reaction. Fast Red (Sigma-Aldrich, catalog # N3020) was used for counterstaining. Slides were allowed to dry and coverslipped using DPX mounting reagent (SDFCL, catalog #46029).

### Image acquisition and analysis

Bright-field images were acquired using Zeiss Axioskop-2 plus microscope equipped with a Nikon DS-fi2 camera and associated NIS Elements V4.0 software. Mouse sections were imaged in Olympus FluoView 1200 confocal microscope with FluoView software. Image analysis was done with Fiji ImageJ software, and/or Adobe Photoshop CS6. Image stitching (for Figure 4G) was performed using the “pairwise stitching” plugin in Fiji. For all stitching operations “subpixel accuracy” parameter was selected. Nonlinear operation such as gamma correction was not performed in any of the figures. Brightness and contrast adjustments were performed identically for control and mutant conditions. For Fig. 4C, cell counting was performed using the “cell counter” plugin in Fiji. All schematics were prepared using Microsoft PowerPoint 2021.

### qPCR Analysis

Hippocampus samples from control and Lmx1aCre; β-Catenin GOF E14.5 embryos were collected in a 1.5 ml tube in 200μl of Trizol® reagent (2 brains, 4 hippocampi per biological replicate). RNA was extracted following the manufacturer’s protocol. RNA concentration was measured using RNA-HS assay kit (catalog #Q32852) in Qubit-2 fluorometer (catalog #Q32866). cDNA was synthesized using SuperScript™ IV kit (catalog #18091050). Real-Time qPCR reactions were performed in triplicates (technical replicates) using KAPA SYBR FAST qPCR Kit (2X) (catalog #KK4601) on LightCycler^®^ 96 Real-time PCR system following the manufacturer’s recommendation. Annealing temperatures and primer concentrations were optimized using gradient PCR for every primer. Melt curve analysis was performed to rule out the possibility of nonspecific amplification. 18S ribosomal RNA was used as a reference gene, and analysis was performed using the ΔΔ threshold cycle (Ct) method. The fold changes were represented as mean ± SEM.

### Primer sequences

*Reelin* (FP: CCACCAAATTTTGTCTCAGGC, RP: AGAAAACTCCAAGCTGACGC),

*Trp73* (FP: CCTTCACTTGCTCACCCTCT, RP: AGTGGAATCGACCTGTACGT),

*Lhx5* (FP: GCAAAACCAACCTCTCGGAG, RP: CTCCGGTGGATAGCTGCTTA),

*Ppp2r2c* (FP: TGGAGTTGGCTTGGATGGTA, RP: ACCAGCTTGACTTTCCCTGA)

### TUNEL Assay

Terminal deoxynucleotidyl transferase dUTP nick end labeling (TUNEL) was performed on E12.5 sagittal sections using the Click-iT™ Plus TUNEL Assay Kit (Cat. No. C10617; Thermo Fisher Scientific) according to the manufacturer’s standard protocol.

### Single-cell RNA sequencing

#### Library preparation

Pregnant dams of control and Lmx1aCre; β-Catenin genotypes were sacrificed on embryonic days 11.5 and 14.5, and embryos were dissected out in ice-cold PBS. They were screened under an epifluorescence lamp for Ai9 (tdTomato) signal and the midline tissue was dissected out with #5 forceps. The tissue was manually dissociated according to the Miltenyi Neural Kit Dissociation Kit (P) protocol. 5 embryos per library were used for the E11.5 time point, and 3 embryos per library were used for the E14.5 time point. Only male embryos were used for the experiment, since the Lmx1aCre transgene is present on the X chromosome and shows a mosaic expression due to random X-inactivation in the females. 3 libraries (E11.5 control, E14.5 control, E14.5 mutant) were prepared according to the instructions of the manufacturer (10X Genomics Chromium Next GEM Single Cell 3’ Gem Kit v3.1). Libraries were sequenced on an Illumina Novaseq 6000 platform using an SP100 flow cell.

#### Data analysis

The FASTQ files were aligned via <u>CellRanger</u> (v7.0), with default parameters, to a custom reference genome, created by adding a 2399 bp tdTomato sequence to mm10 for the purpose of tracking Ai9 reporter expression. Filtered feature barcode matrices were processed for each library individually using Seurat v5. Preprocessing involved filtering cells based on calculated mitochondrial gene percentage and number of cells and features, to keep only those within three median absolute deviations of three population medians (+-3 MAD). The resulting gene expression matrices were log normalised and the top 2000 variable features were calculated. The expression of all genes were scaled across all cells and PCA was run with the variable features. The top 20 PCs were used to find neighbours and create UMAPs. Marker genes from literature were used to annotate known cell types.

The E11.5 and E14.5 control libraries were merged, then jointly log-normalized and scaled across all cells. Harmony was used for integration. The clusters which showed tdTomato expression were subset. Monocle3 was used with default parameters to obtain pseudotime trajectories. A similar workflow was followed to integrate E14.5 control and β-Catenin GOF libraries, but Harmony was not used for integration, since we wanted to avoid artificial integration of any cell types that were biologically different. To avoid batch effects as much as possible, we sacrificed timed pregnant mice of both genotypes at E14.5 on the same day, and every downstream step from tissue dissociation to microfluidic droplet partitioning and library preparation was done at the same time with the same batch of reagents in parallel wells. No regression was performed according to genotype.

At E14.5, differentially expressed genes were calculated between control and β-Catenin GOF neurons using Seurat’s FindMarkers() function using the Wilcoxon Rank Sum test. Genes were considered significant if they exhibited an adjusted p-value < 0.05 (Bonferroni correction) and a natural log fold-change > 0.1, with a minimum detection threshold of 1% of cells in either cluster (min.pct = 0.1). The top 20 genes were used to create a control and β-Catenin GOF module. Seurat’s AddModuleScore function was used (with default parameters) to calculate the average expression levels of each gene in a user-defined “module” on single cell level, subtracted by the aggregated expression of control genes randomly selected from each bin. Cells with a module score above the 99th percentile in the entire dataset were highlighted.

### Statistics and reproducibility

Biological replicates (n) denote samples that have been obtained from individual embryos/pups. Blinded analysis was not possible for Lmx1aCre genotypes, since they are easily distinguishable by phenotypic features. Blinded analysis was performed for Foxj1Cre animals. Statistical analyses were performed in GraphPad Prism (v9.3.1) and in R (v4.4.0). Information about statistical tests and p-values are provided in corresponding figure legends. Distribution of data points was analysed using the Shapiro-Wilk normality test. If the data is distributed normally, then a parametric test was chosen, otherwise a non-parametric test was performed. For all statistical tests, the chosen confidence interval is 95% (α=0.05). For the XY plots in Figure 1C, Figure 2D, Figure 4C, Figure 4F error bars represent SEM.

## Data Availability

The data generated in this study is available at the NCBI SRA (Accession number PRJNA1141348) and NCBI GEO (Accession number GSE273962) databases.

## Acknowledgements

We thank the following for kindly gifting us mouse lines used in this study: Kathleen J. Millen (Seattle Children’s Hospital) for Lmx1aCre; Makoto M. Taketo (Kyoto University) for Ctnnb1 fl/fl; Michael. J. Holtzman and Yong Zhang (University of Washington, St. Louis) for Foxj1Cre. We thank Shital Suryavanshi and the animal house staff of the Tata Institute of Fundamental Research (TIFR), GH Mohan and the staff at National Centre for Biological Sciences (NCBS) for their invaluable support. We thank Santosh Srirangam (10X Genomics), Awadhesh Pandit and Lakshminarayanan C P (NCBS-TIFR NGS facility) for experimental design consultation and library preparation for the single-cell sequencing experiments. We thank Vishal Nanavaty (Neuberg Center for Genomic Medicine, Neuberg Supratech Reference Laboratory) for providing 10X Genomics reagents and sequencing services.

## CRediT Authorship Contribution Statement

Conceptualization: AS, AP, ST; Data Curation: AS, AP; Formal analysis: AS, AP; Funding acquisition: ST, AP; Investigation: AS, AP, DD, MC; Methodology: AS, AP; Project administration: ST; Resources: ST; Supervision: ST; Validation: AS, AP, DD; Visualization: AS, AP; Writing - original draft: AS, ST; Writing - review & editing: AS, ST

