## Supplementary Information for "Cajal-Retzius fate specification is disrupted by constitutive activation of β-Catenin in hem progenitors"

Figure S1

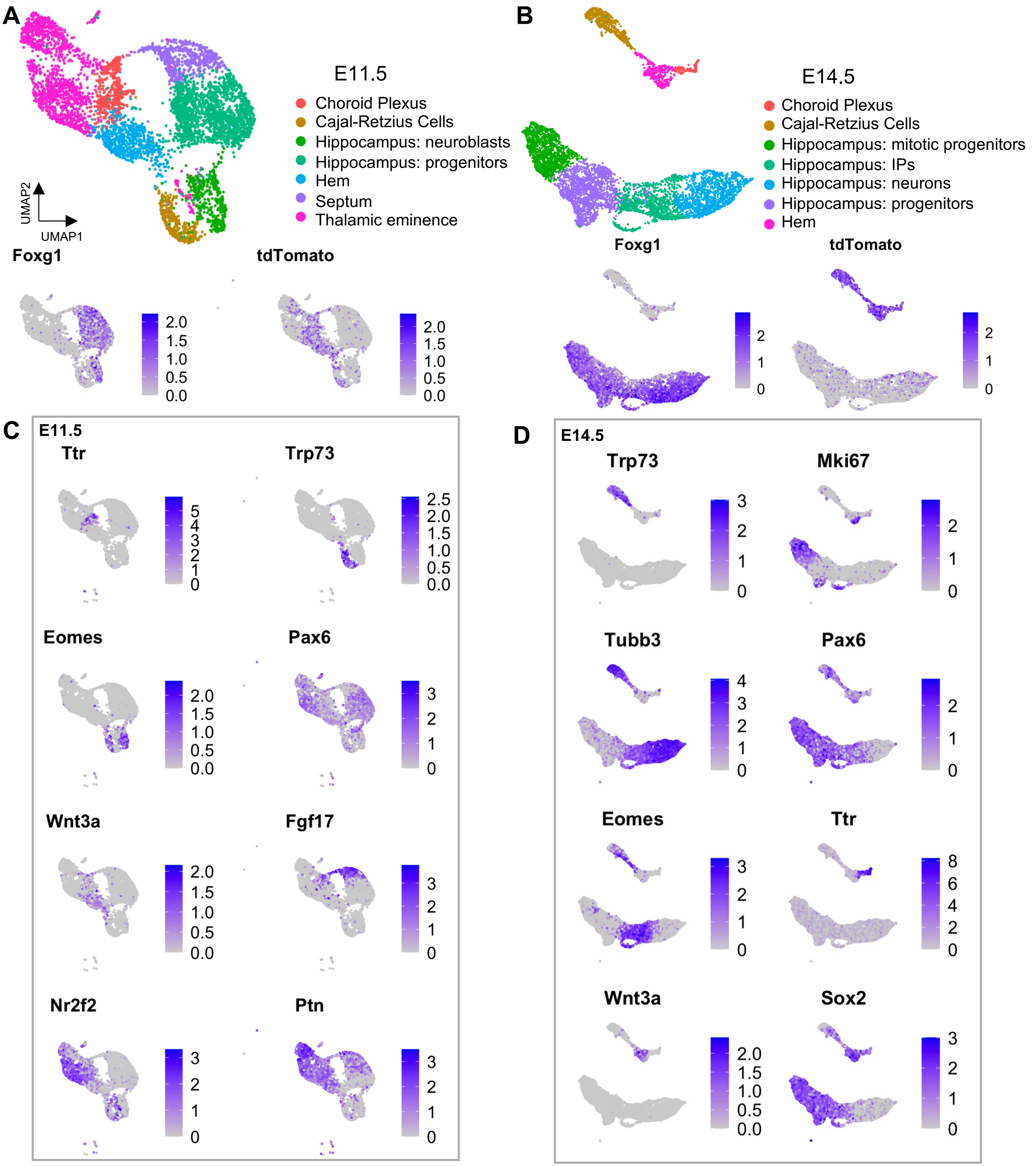

A
B
E11.5
Choroid Plexus
Cajal-Retzius Cells
Hippocampus: neuroblasts
Hippocampus: progenitors
Hem
Septum
Thalamic eminence
UMAP2
UMAP1
E14.5
Choroid Plexus
Cajal-Retzius Cells
Hippocampus: mitotic progenitors
Hippocampus: IPs
Hippocampus: neurons
Hippocampus: progenitors
Hem
Foxg1
tdTomato
C
E11.5
Ttr
Trp73
Eomes
Pax6
Wnt3a
Fgf17
Nr2f2
Ptn
D
E14.5
Trp73
Mki67
Tubb3
Pax6
Eomes
Ttr
Wnt3a
Sox2

**Figure S1. Annotation of the E11.5 and E14.5 scRNA-seq libraries.**

(A, B) UMAP representing the entire dorsal midline cell population isolated for scRNAseq at E11.5 (A) and 14.5 (B).

(C,D) Expression of cell type enriched genes at each age.

#### Figure S2

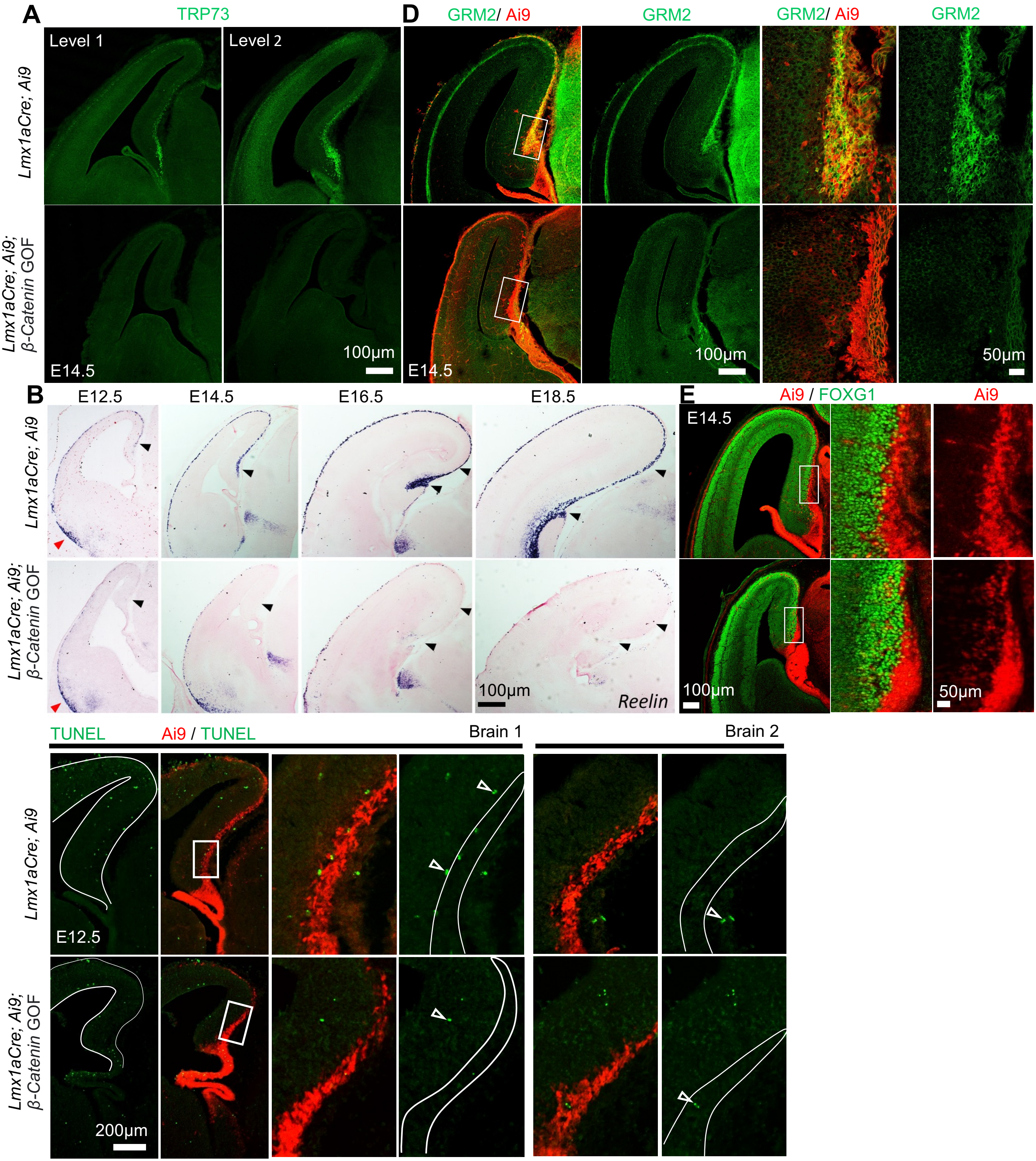

A
TRP73
Level 1
Level 2
Lmx1aCre; Ai9
Lmx1aCre; Ai9; β-Catenin GOF
E14.5
100µm
D
GRM2/ Ai9
GRM2
GRM2/ Ai9
GRM2
E14.5
100µm
50µm
B
E12.5
E14.5
E16.5
E18.5
Lmx1aCre; Ai9
Lmx1aCre; Ai9; β-Catenin GOF
100µm
Reelin
E
Ai9 / FOXG1
Ai9
E14.5
100µm
50µm
TUNEL
Ai9 / TUNEL
Brain 1
Brain 2
Lmx1aCre; Ai9
Lmx1aCre; Ai9; β-Catenin GOF
E12.5
200µm

**Figure S2. Characterization of the Ai9+ cells in *Lmx1aCre; β-Catenin* GOF brains.**

(A) TRP73 in control and *Lmx1aCre; β-Catenin* GOF midline at E14.5 shows no apparent signal in the GOF.

(B) *Reelin* expression across developmental stages in control and *Lmx1aCre; β-Catenin* GOF brains shows no apparent signal in the cortical marginal zone (black arrowheads), while signal is visible near the PSPB (red arrowheads).

(C) Ai9 labels the cortical hem and its derivatives in E12.5 control and *Lmx1aCre; β-Catenin* GOF brains. TUNEL staining shows no increase in apoptotic Ai9+ cells in the marginal zone, in three replicates.

(D) GRM2 in control and *Lmx1aCre; β-Catenin* GOF midline at E14.5 shows no apparent signal in the GOF.

(E) FOXG1 in control and *Lmx1aCre; β-Catenin* GOF midline at E14.5 shows no signal in Ai9+ cells in both conditions.

#### Figure S3

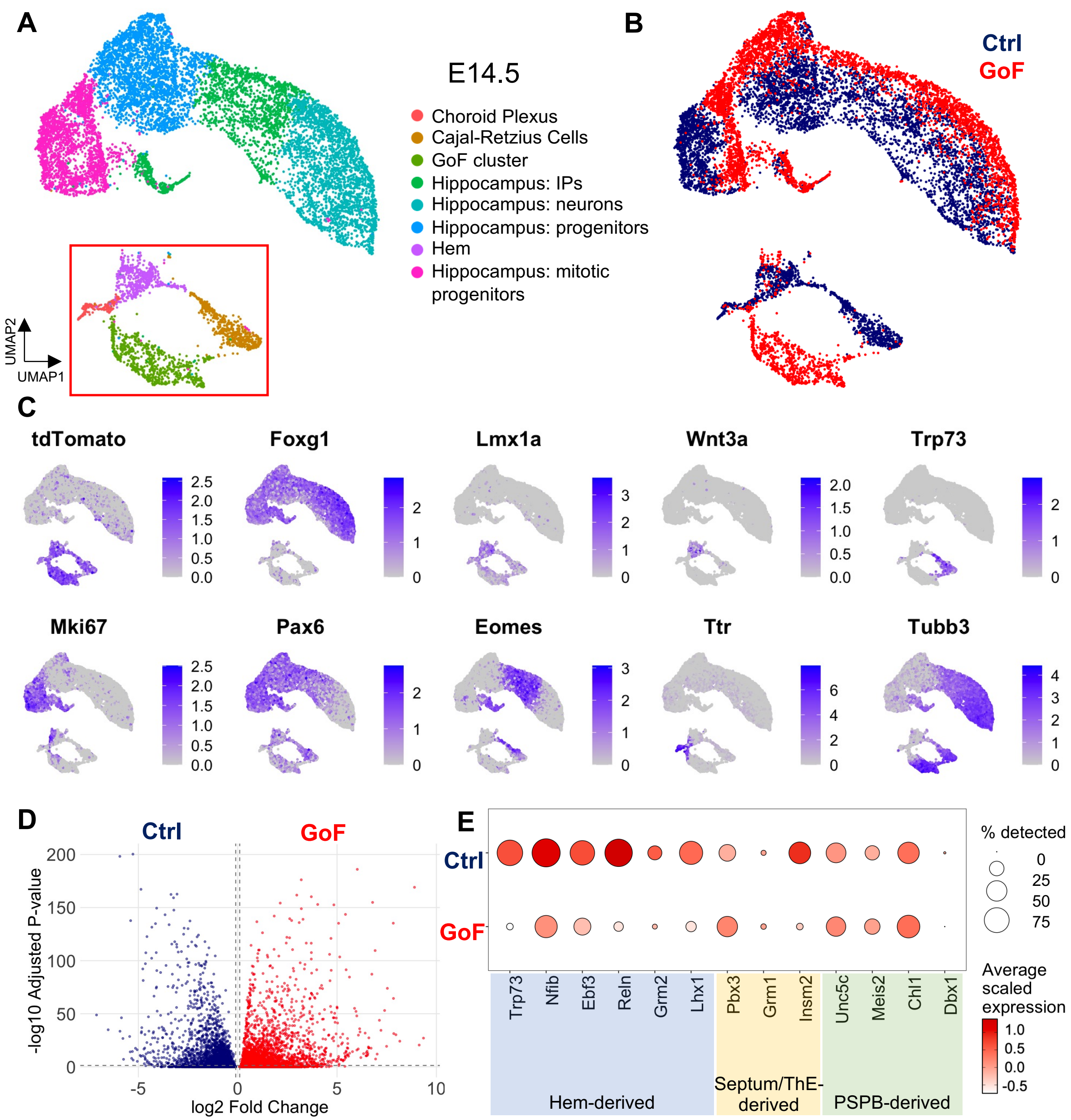

A
E14.5
Choroid Plexus
Cajal-Retzius Cells
GoF cluster
Hippocampus: IPs
Hippocampus: neurons
Hippocampus: progenitors
Hem
Hippocampus: mitotic progenitors
UMAP2
UMAP1
B
Ctrl
GoF
C
tdTomato
Foxg1
Lmx1a
Wnt3a
Trp73
Mki67
Pax6
Eomes
Ttr
Tubb3
D
Ctrl
GoF
-log10 Adjusted P-value
log2 Fold Change
E
Ctrl
GoF
Trp73
Nfib
Ebf3
Reln
Grm2
Lhx1
Pbx3
Grm1
Insm2
Unc5c
Meis2
Chl1
Dbx1
Hem-derived
Septum/ThE-derived
PSPB-derived
% detected
Average scaled expression

**Figure S3. Merged scRNAseq libraries from control and *Lmx1aCre; β-Catenin* GOF midline**

(A, B) UMAPs showing the entire integrated control and *Lmx1aCre; β-Catenin* GOF dataset from E14.5, colour-coded by genotype

(C) Expression of cell type specific enriched genes in UMAPs from A, B.

(D) Volcano plot showing the distribution of genes upregulated in control CR cells and *Lmx1aCre; β-Catenin* GOF neurons.

(E) Dot plots showing scaled expression levels of marker genes for different subtypes of CR cells in control and *β-Catenin* GOF neurons.

### Supplementary Table 1. Modules for Control (CR) and GoF (hem-derived) neurons

| **Control (CR) module** | **GoF (hem-derived) module** |
|---|---|
| *Trp73* | *Wif1* |
| *Reln* | *Rpl29* |
| *Tbr1* | *Sostdc1* |
| *Cacna2d1* | *Tenm2* |
| *Fat3* | *Nkd1* |
| *Nnat* | *Mybpc1* |
| *Agbl4* | *Nap1l1* |
| *Slc17a6* | *Apcdd1* |
| *Nrn1* | *Rbfox1* |
| *Palmd* | *Sp5* |
| *Elavl4* | *Phlda1* |
| *Lhx9* | *Bsg* |
| *Plcl1* | *Zic4* |
| *Tmem158* | *Wwc1* |
| *Tanc2* | *Axin2* |
| *Cacna2d2* | *Tpt1* |
| *Maml13* | *Igfbp2* |
| *Ripor2* | *Notum* |
| *Nfix* | *Asic2* |
| *Nfib* | *Rpl7* |
